# Seven-CpG DNA Methylation Age determined by Single Nucleotide Primer Extension and Illumina’s Infinium MethylationEPIC array provide highly comparable results

**DOI:** 10.1101/2021.08.13.456213

**Authors:** Valentin Max Vetter, Christian Humberto Kalies, Yasmine Sommerer, Lars Bertram, Ilja Demuth

**Affiliations:** Charité – Universitätsmedizin Berlin, corporate member of Freie Universität Berlin, Humboldt-Universität zu Berlin, and Berlin Institute of Health, Lipid Clinic at the Interdisciplinary Metabolism Center, Germany; Department for Psychology, Humboldt University Berlin, Berlin, Germany; Lübeck Interdisciplinary Platform for Genome Analytics (LIGA), University of Lübeck, Lübeck, Germany; Center for Lifespan Changes in Brain and Cognition (LCBC), Dept of Psychology, University of Oslo, Oslo, Norway; Charité - Universitätsmedizin Berlin, BCRT - Berlin Institute of Health Center for Regenerative Therapies, Berlin, Germany

**Author notes:** **Corresponding author:** Ilja Demuth (Ph.D.), Charité - Universitätsmedizin Berlin, Lipid Clinic at the Interdisciplinary Metabolism Center, Biology of Aging Group, Augustenburger Platz 1, 13353 Berlin.

**Keywords:** Epigenetic Clock, DNA Methylation Age, Biological Age, Aging, Single-Nucleotide Primer Extension

## Abstract

DNA methylation age (DNAm age, epigenetic clock) is a novel and promising biomarker of aging. It is calculated from the methylation fraction of specific cytosine phosphate guanine sites (CpG sites) of genomic DNA. Several groups have proposed epigenetic clock algorithms and these differ mostly regarding the number and location of the CpG sites considered and the method used to assess the methylation status. Most epigenetic clocks are based on a large number of CpGs, e.g. as measured by DNAm microarrays. We have recently evaluated an epigenetic clock based on the methylation fraction of seven CpGs that were determined by methylation-sensitive single nucleotide primer extension (MS-SNuPE). This method is more cost-effective when compared to array-based technologies as only a few CpGs need to be examined. However, there is only little data on the correspondence in epigenetic age estimation using the 7-CpG clock and other algorithms.

To bridge this gap, in this study we measured the 7-CpG DNAm age using two methods, via MS-SNuPE and via the MethylationEPIC array, in a sample of 1,058 participants of the Berlin Aging Study II (BASE-II), assessed as part of the GendAge study. On average, participants were 75.6 years old (SD: 3.7, age range: 64.9 – 90.0, 52.6% female). Agreement between methods was assessed by Bland-Altman plots. DNAm age was highly correlated between methods (Pearson’s r=0.9) and Bland-Altman plots showed a difference of 3.1 years. DNAm age by the 7-CpG formula was 71.2 years (SD: 6.9 years, SNuPE) and 68.1 years (SD: 6.4 years, EPIC array). The mean of difference in methylation fraction between methods for the seven individual CpG sites was between 0.7 and 13 percent. To allow direct conversion between methods we developed an adjustment formula with a randomly selected training set of 529 participants using linear regression. After conversion of the Illumina data in a second and independent validation set, the adjusted DNAm age was 71.44 years (SD: 6.1 years, n=529). In summary, we found the results of DNAm clocks to be highly comparable. Furthermore, we developed an adjustment formula that allows for direct conversion of estimates between methods and enables one singular clock to be used in studies that employ either the Illumina or the SNuPE method.

## Introduction

DNA methylation age (epigenetic clock, DNAm age) and its deviation from chronological age, DNAm age acceleration (DNAmAA), are novel and intensively studied biomarkers of biological aging. They are calculated from the methylation fraction of specific cytosine phosphate guanine (CpG) sites of genomic DNA. Numerous epigenetic clocks are available that differ in location and number of analyzed CpG sites and how these sites were selected. Their associations with mortality (reviewed in [1] and meta-analysis [2]), morbidity and age associated phenotypes (reviewed in [3]) are well documented.

The most frequently used clocks include 71 [4] or more CpG sites [5, 6] and therefore have to rely on epigenome-wide measurements, mostly carried out by Illumina’s array-based “Infinium Methylation Assays”. As an addition to these “big” epigenetic clocks, we recently reported a novel 7-CpG epigenetic clock whose underlying methylation data were measured by the methylation-sensitive nucleotide primer extension method (MS-SNuPE) [7]. This methodological approach was originally described by Kaminsky et al. [8] and modified for the calculation of methylation age by Vidal-Bralo and colleagues [9, 10]. When the interest is only seven CpGs this method is more cost effective compared to methylation arrays producing genome-wide methylation data. Consequently, the SNuPE method has been used in several studies [9, 11, 12]. However, the comparability of the findings described in these reports with reports relying on the identical CpG sites determined by an array-based genome-wide (e.g. Liu et al. [13]) approach is uncertain.

This study aims to close this gap by measuring the methylation fractions of the regarding CpG sites with both the MS-SNuPE method and the “Infinium MethylationEPIC” array (Illumina Inc.) in a cohort of 1,058 adults (female: 52.6%). We subsequently compared the 7-CpG DNAm age of both methods. Due to the strong linear association of the results generated by both methods, we propose an adjustment formula that allows for direct conversion between 7-CpG methylation age measured with the EPIC array and the SNuPE method.

## Materials and Methods

### BASE-II/GendAge Study

The multi-disciplinary and longitudinal BASE-II study aims to identify factors that are associated with “healthy vs. unhealthy” aging [14, 15]. The medical follow-up assessments took place between 2018 and 2020 and were part of the GendAge study [16]. The current study included 1,058 GendAge participants with a mean age of 75.6 years (SD: 3.7 years, age range: 64.9 – 90.0 years, 52.6 % female). For detailed information on BASE-II and GendAge please refer to Bertram et al. [14] and Demuth et al. [16].

The GendAge study was executed in accordance with the Declaration of Helsinki and approved by the ethics committee of the Charité – Universitätsmedizin Berlin (approval number: EA2/144/16). All participants gave written informed consent and GendAge is registered in the German Clinical Trials registry (DRKS00016157).

### Seven-CpG epigenetic clock

The epigenetic clock used in this study employs seven CpG sites: cg09809672, cg02228185, cg19761273, cg16386080, cg17471102, cg24768561 and cg25809905. The formula was trained with SNuPE methylation data obtained from participants at baseline examination of the BASE-II [7] and is referred to as “BII7”-formula. The described CpG sites were previously identified to be the most informative on chronological age and to be measurable in a SNuPE assay by Vidal-Bralo and colleagues [9, 10]. An additional CpG site (cg10917602) was measured as well but is not included in the 7-CpG clock. Its results are shown in Supplementary Figure 1 to 3.

### DNA Methylation assessment using: Methylation-Sensitive Single Nucleotide Primer Extension (MS-SNuPE)

The analyzed genomic DNA was extracted from EDTA whole blood samples with the LGC “Plus XL manual kit”, LGC, UK, and stored at -20°C. Briefly, 1000 ng genomic DNA were bisulfite converted with the “EZ-96 DNA Methylation-Lightning Kit”, Zymo Research. Subsequently, a multiplex PCR was conducted to amplify DNA sections surrounding the CpG sites of interest. The sample was cleaned from remaining oligonucleotides and dNTPs with “Shrimp Alkaline Phosphatase”, Affymetrix, and “Exonuclease I”, New England Biolabs. The “SNaPshot Multiplex Kit”, Applied Biosystems, was used for the single nucleotide primer extension (SNuPE). After an ultimate cleaning step with “Shrimp Alkaline Phosphatase”, 6 µg of the SNuPE-products were measured with an “3730 DNA Analyzer”, Applied Biosystems and HITACHI. Raw data was inspected and processed with the “GeneMapper” software package, Applied Biosystems. The peak height was used to calculate the individual methylation fraction. For a more detailed description of the MS-SNuPE protocol, please refer to reference [7].

Extrinsic epigenetic age acceleration (EEAA) was calculated as residuals of a linear regression analysis of DNAm age on chronological age. Intrinsic epigenetic age acceleration (IEAA) was calculated as residuals of a linear regression analysis of DNAm age on chronological age and cell counts of neutrophils, monocytes, lymphocytes and eosinophiles. The latter is a modified version of the IEAA proposed by Quach and colleagues [17].

### DNA Methylation assessment using: Infinium MethylationEPIC array

DNA methylation data was additionally obtained from the same DNA samples with the “Infinium MethylationEPIC” array by Illumina. Recommended default parameters were used for data pre-processing with the R-package “Bigmelon” [18] and R 3.6.1 [19].

Briefly, probes were removed from the analyses if they had 1% or more samples with a detection p-value of 0.05 or a bead count below three in more than 5% of the samples. Outliers were identified by the *outlyx* and *pcout* [20] function. Bisulfite conversion efficiency was estimated by *bscon* and samples with values <80% were excluded from all following analyses. Subsequently, the samples were reloaded without the identified outliers and the function *dasen* was used for normalization. The amount of change in beta values due to normalization was determined by the function *qual*. Samples with a root-mean-square deviation of ≥0.1 in beta value after normalization were removed and loading and normalization were repeated with the new sample set. DNAmAA with the Illumina data was calculated as EEAA and IEAA in the same way as described above.

### Statistical analysis

The statistical analyses and figures presented in this study were executed and designed with R 3.6.2 [19] and the “ggplot2” package [21]. Bland-Altman plots and statistics were computed with the “blandr” package [22]. Linear Regression analyses were calculated with R’s “lm” function. Methylation data of both methods was available in 1,087 of the older participants. Participants with difference between measurement methods of 3 SD or more were excluded from all analyses (n=29). For the purpose of developing an adjustment formula converting the DNAm age from EPIC data into SNuPE data, we split the full BASE-II dataset as assessed in GendAge into a “training” and “validation” subsample. Individuals in either subset were randomly selected with the “sample” function in R. Throughout the study, statistical significance was assumed for p-values<0.05.

## Results

### GendAge study cohort characteristics

We analyzed 1,058 participants of the GendAge study which included the medical follow-up assessments of BASE-II participants. The chronological age of the participants ranged between 64.9 and 90.0 years and was on average 75.6 years (SD: 3.7) and 52.6% of the analyzed participants were female. Cohort characteristics are displayed in Table 1.

**Table 1:**
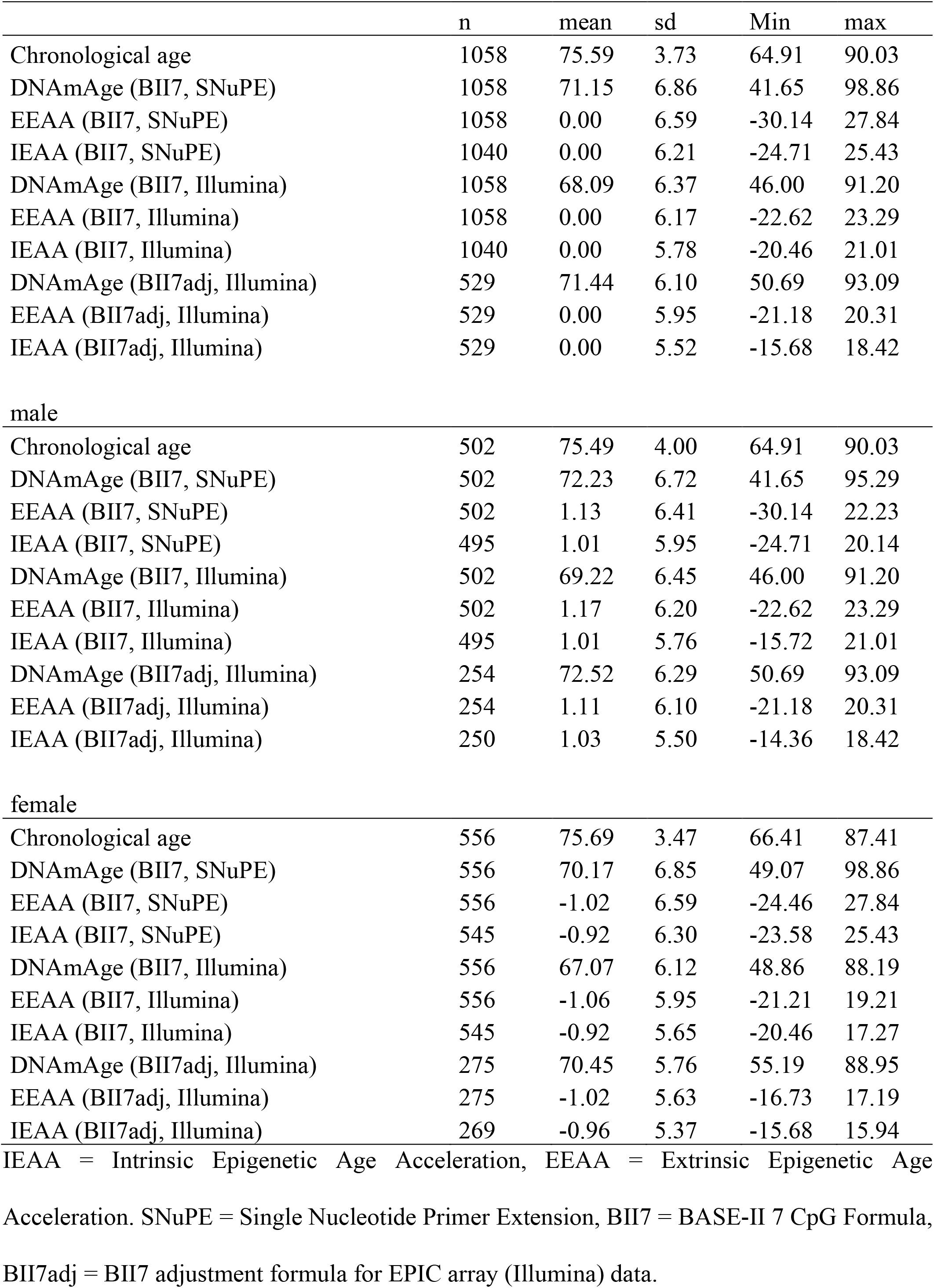
Cohort characteristics

|  | n | mean | sd | Min | max |
| --- | --- | --- | --- | --- | --- |
| Chronological age | 1058 | 75.59 | 3.73 | 64.91 | 90.03 |
| DNAmAge (BII7, SNUPE) | 1058 | 71.15 | 6.86 | 41.65 | 98.86 |
| EEAA (BII7, SNUPE) | 1058 | 0.00 | 6.59 | -30.14 | 27.84 |
| IEAA (BII7, SNUPE) | 1040 | 0.00 | 6.21 | -24.71 | 25.43 |
| DNAmAge (BII7, Illumina) | 1058 | 68.09 | 6.37 | 46.00 | 91.20 |
| EEAA (BII7, Illumina) | 1058 | 0.00 | 6.17 | -22.62 | 23.29 |
| IEAA (BII7, Illumina) | 1040 | 0.00 | 5.78 | -20.46 | 21.01 |
| DNAmAge (BII7adj, Illumina) | 529 | 71.44 | 6.10 | 50.69 | 93.09 |
| EEAA (BII7adj, Illumina) | 529 | 0.00 | 5.95 | -21.18 | 20.31 |
| IEAA (BII7adj, Illumina) | 529 | 0.00 | 5.52 | -15.68 | 18.42 |
| male |  |  |  |  |  |
| Chronological age | 502 | 75.49 | 4.00 | 64.91 | 90.03 |
| DNAmAge (BII7, SNUPE) | 502 | 72.23 | 6.72 | 41.65 | 95.29 |
| EEAA (BII7, SNUPE) | 502 | 1.13 | 6.41 | -30.14 | 22.23 |
| IEAA (BII7, SNUPE) | 495 | 1.01 | 5.95 | -24.71 | 20.14 |
| DNAmAge (BII7, Illumina) | 502 | 69.22 | 6.45 | 46.00 | 91.20 |
| EEAA (BII7, Illumina) | 502 | 1.17 | 6.20 | -22.62 | 23.29 |
| IEAA (BII7, Illumina) | 495 | 1.01 | 5.76 | -15.72 | 21.01 |
| DNAmAge (BII7adj, Illumina) | 254 | 72.52 | 6.29 | 50.69 | 93.09 |
| EEAA (BII7adj, Illumina) | 254 | 1.11 | 6.10 | -21.18 | 20.31 |
| IEAA (BII7adj, Illumina) | 250 | 1.03 | 5.50 | -14.36 | 18.42 |
| female |  |  |  |  |  |
| Chronological age | 556 | 75.69 | 3.47 | 66.41 | 87.41 |
| DNAmAge (BII7, SNUPE) | 556 | 70.17 | 6.85 | 49.07 | 98.86 |
| EEAA (BII7, SNUPE) | 556 | -1.02 | 6.59 | -24.46 | 27.84 |
| IEAA (BII7, SNUPE) | 545 | -0.92 | 6.30 | -23.58 | 25.43 |
| DNAmAge (BII7, Illumina) | 556 | 67.07 | 6.12 | 48.86 | 88.19 |
| EEAA (BII7, Illumina) | 556 | -1.06 | 5.95 | -21.21 | 19.21 |
| IEAA (BII7, Illumina) | 545 | -0.92 | 5.65 | -20.46 | 17.27 |
| DNAmAge (BII7adj, Illumina) | 275 | 70.45 | 5.76 | 55.19 | 88.95 |
| EEAA (BII7adj, Illumina) | 275 | -1.02 | 5.63 | -16.73 | 17.19 |
| IEAA (BII7adj, Illumina) | 269 | -0.96 | 5.37 | -15.68 | 15.94 |

IEAA = Intrinsic Epigenetic Age Acceleration, EEAA = Extrinsic Epigenetic Age
Acceleration. SNUPE = Single Nucleotide Primer Extension, BII7 = BASE-II 7 CpG Formula,
BII7adj = BII7 adjustment formula for EPIC array (Illumina) data.

### Comparison of DNA methylation fraction measured by the SNuPE method and Illumina’s EPIC array

We measured the methylation fraction of the sites included in the 7-CpG clock (cg09809672, cg02228185, cg19761273, cg16386080, cg17471102, cg24768561 and cg25809905) in the same DNA samples with the SNuPE method and with Illumina’s “Infinium MethylationEPIC” array. The methylation fraction of cg1097602 was assessed as well but is not included in the 7- CpG clock. The results are shown in Supplementary Figure 1 to 3.

Participants showed methylation fractions between 4.7 % (cg19761273) and 88% (cg25809905) in SNuPE data and between 9.7 % (cg19761273) and 81.5% (cg02228185) in Illumina data (Figure 1 A). The average methylation range per CpG site was 44.9 percentage points (SNuPE) and 39.1 percentage points (Illumina).

**Figure 1:**
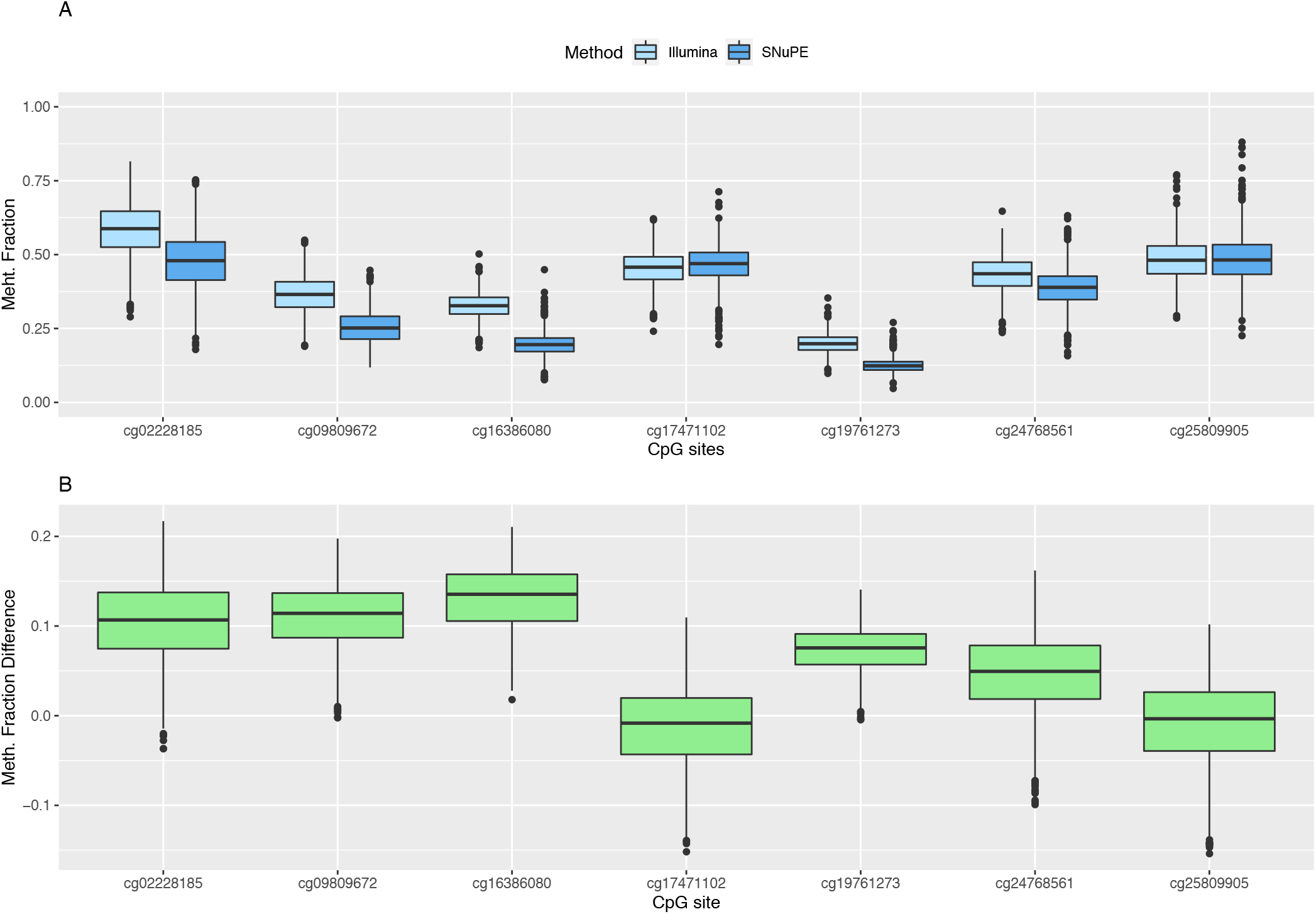
Boxplots of methylation fraction measured by the SNuPE and EPIC array method (A) and difference between both methods (B).

The mean of the differences between methods ranged between 0.7 (cg25809905) and 13.1 (cg16386080) percentage points (Figure 1 B). The smallest difference was found in CpG sites whose mean methylation fraction was close to 50%. The methylation fractions of individual CpG sites between methods were moderately (cg16386080, Pearson’s r=0.62) to highly correlated (cg02228185, Pearson’s r=0.89) (Figure 2, A-H).

**Figure 2:**
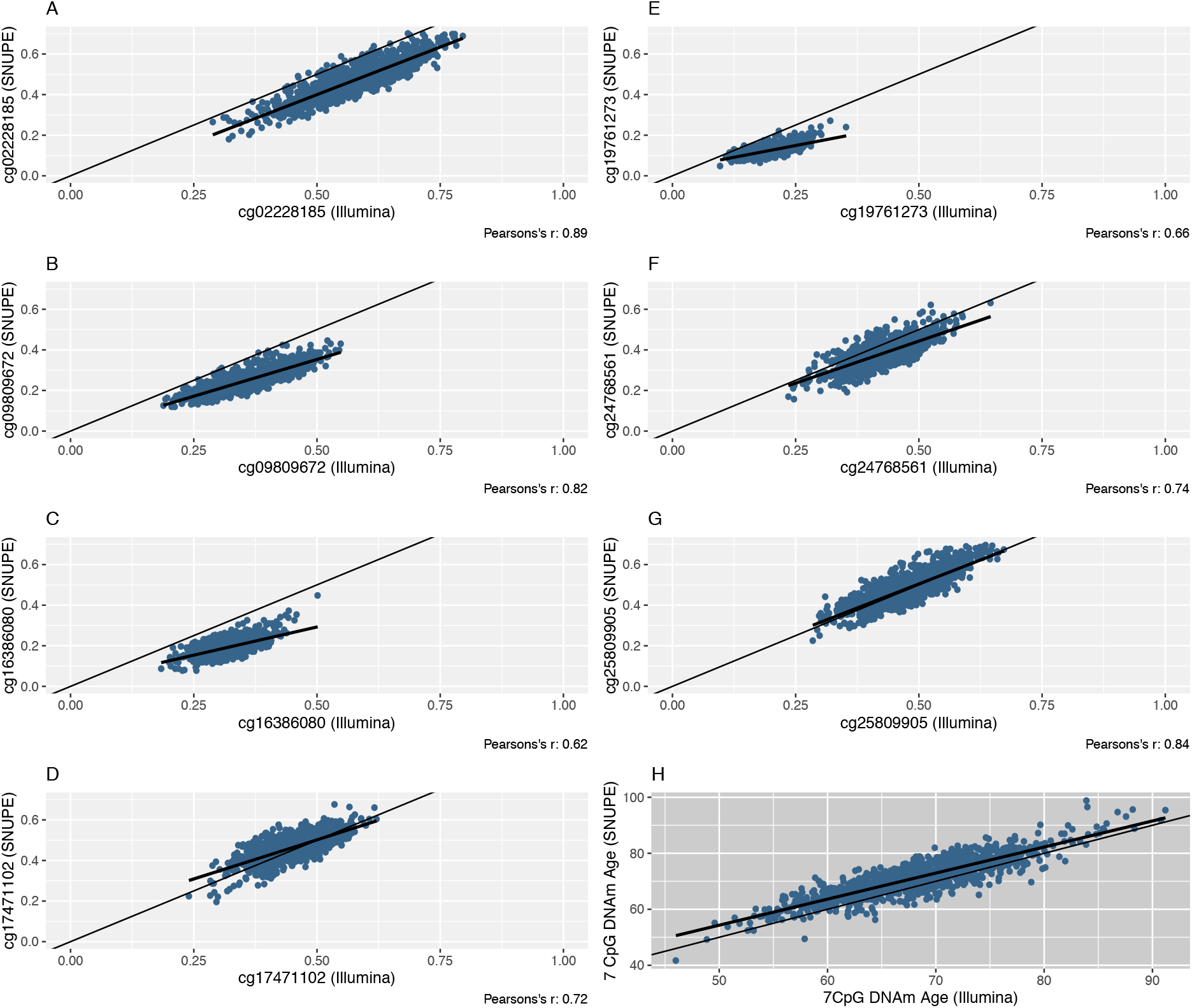
Scatterplots of the methylation fraction of seven CpG sites that were measured with the SNuPE and EPIC array (Illumina) method (A-G). The line of equality (thin) and the regression line (bold) are displayed. The DNAm age, that was calculated with the 7 CpG clock (“BII7” formula) is shown in scatterplot H.

To assess the limits of agreement and investigate a possible association between the measurement error and the methylation fraction, Bland-Altman plots were computed (Supplementary Figure 1). A regression analysis of the parameters analyzed in the Bland-Altman plots was generated to objectify a potential proportional bias. All plots showed marginal skewness at most (ß ≤ |0.19|), except for one CpG site that showed a weak negative association (ß=-0.42, cg19761273) between the difference of measurements and the mean of measurements. The latter was proposed by Bland and Altman as a substitute for the unknown true value [23].

We found a mean difference between measurement methods of 7 percentage points or less in cg25809905, cg17471102, cg24768561 and cg19761273 (Figure 1B). The highest average difference in the methylation fraction between methods was 13 percentage points and was found in cg16386080. The limits of agreement (LOA), which were pinned by Bland and Altman [23] as mean of difference +/- 1.96 SD, contain by definition 95% of the values that represent the differences between methods. The range between the upper LOA (mean of difference + 1.96SD) and the lower LOA (mean of difference – 1.96SD) was 18.5 percentage points or smaller.

### Seven-CpG DNAm Age is highly correlated between methods

The DNAm age was calculated with the 7-CpG formula (“BII7”) that was trained on SNuPE methylation data [7]. It was measured to be on average 71.2 years by the SNuPE method and 68.1 years by the Illumina method. We found the DNAm age of men to be on average two years higher compared to women although the difference in chronological age was only 0.2 years, a finding that was reported for other epigenetic clocks as well [24, 25].

Both clocks were highly correlated with each other (Pearson’s r=0.86). The deviation of DNAm age around the “line of equality”, indicating the potentially optimal line of identical results in both methods, is shown in Figure 2 H. The Bland-Altman plot (Supplementary Figure 1) shows a consistent bias of 3.1 years with a lower LOA of -3.8 and an upper LOA of 9.9 years. We found no proportional bias (linear regression analysis; ß= 0.08).

### Correlation between DNAm Age and Chronological Age and New Formula for Conversion Between Methods

A moderate correlation was found between chronological age and DNAm age based on SNuPE data (Pearson’s r=0.28) and Illumina data (Pearson’s r=0.25). The slope of the regression line of DNAm age on chronological age was 0.51 (SNuPE) and 0.42 (Illumina) (Figure 3 A). To adjust the DNAm age obtained with EPIC array-based data, we computed a linear regression analysis of SNuPE-DNAm age on Illumina-DNAm age in a training set of 529 randomly selected participants. The resulting adjustment formula (“BII7adj”) allows for direct conversion:

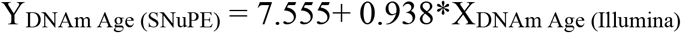

**Figure 3:**
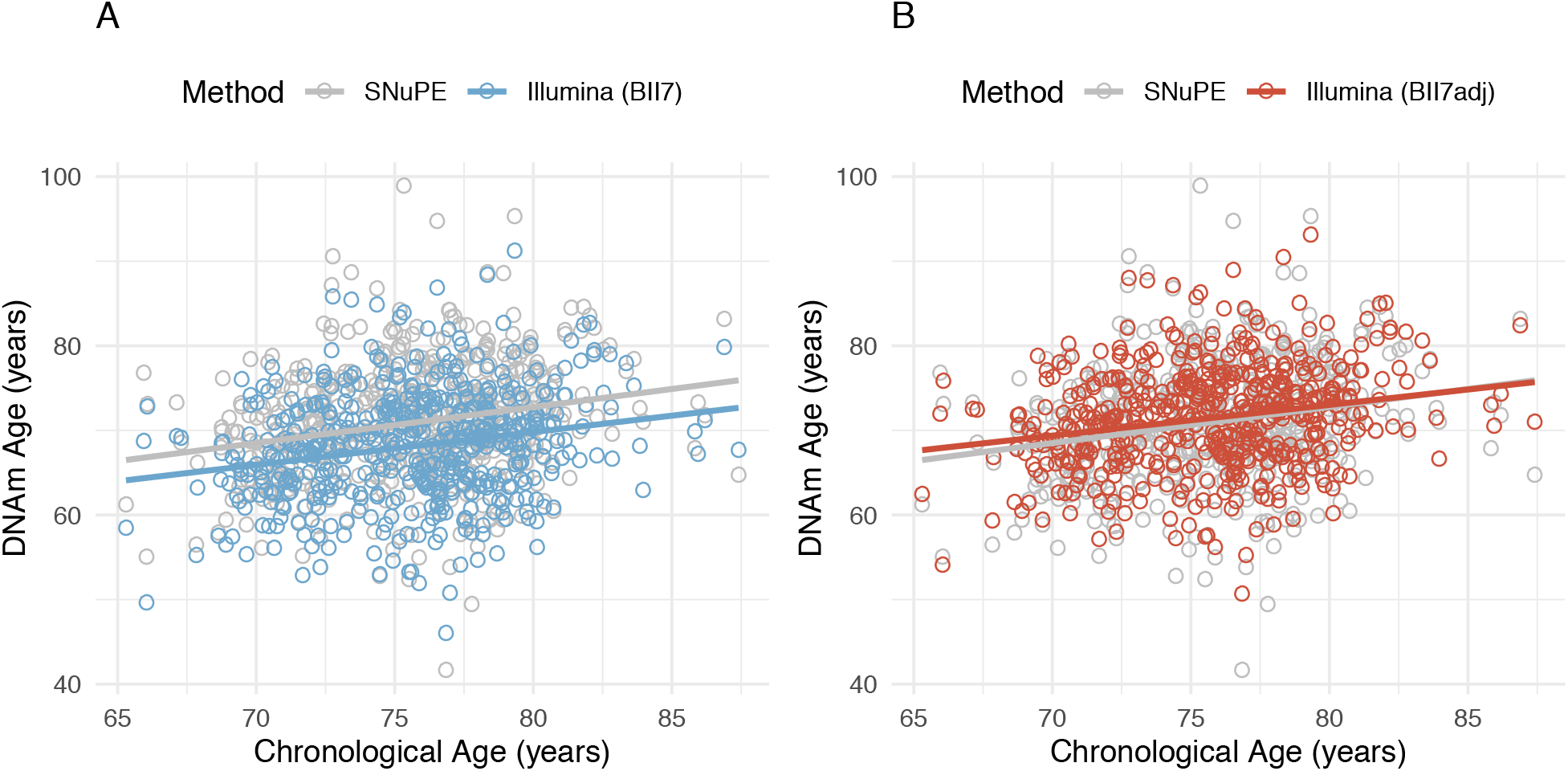
Scatterplot of methylation age, measured with the SNuPE (grey) and EPIC array (Illumina) method (color) calculated with the “BII7” (A) and “BII7adj” formula (B), on chronological age in the validation set (n=529).

Descriptive statistics of the adjusted DNAm age obtained through EPIC array-based data in the validation set of our cohort (n=529) are displayed in Table 1 and its association to chronological age is shown in Figure 3 B.

## Discussion

In this study we compare two methods of DNAm age measurement to construct the 7-CpG DNAm clock, i.e. via the SNuPE method and via high-throughput DNAm profiling using the “InfiniumMethylationEPIC” array, in 1,058 participants of the GendAge study. Although DNAm age estimates strongly correlated between both methods, they showed a deviation of 3.1 years on average. Hence, we propose an adjustment formula to directly convert the results of both methods to increase their comparability. These findings will enable using the 7-CpG clock, that was originally developed for SNuPE data only, to be calculated with methylation data that was obtained with the EPIC array as well. To our knowledge, this is the only study to systematically compare these epigenetic age estimations from the 7-CpG clock using two different methodological approaches in one dataset.

To assess LOA between the methylation measurements of the individual CpG sites, we followed the established and widely used approach that was first proposed by Bland and Altman in 1986 [23]. To evaluate a possible association between the difference between measurements and the true value, which is unknown and therefore estimated as the mean between measurements, Bland-Altman plots were drawn. The range between the upper and lower LOA of the other CpG sites, that includes by definition 95% of the values that represent the differences between measurements, was 18.5 percentage points or smaller, a range which can be accepted for the purpose of DNAm age calculation.

Only one of the epigenetic clocks CpG sites (cg19761273) showed a moderate association between difference between measurements and mean of measurements. However, we found the smallest difference between methods in CpG sites, whose methylation fraction was close to 50%. We therefore assume a potential proportional bias between both methods which, however, (almost) never becomes detectable in our cohort. That may be due to the comparatively small methylation ranges that occur in each individual CpG site in vivo. A series of experiments with DNA whose methylation fraction ranges between 0% and 100% would allow to test for proportional bias in every individual CpG site. Although easy to conduct, this experiment would not influence the evaluation of this method with regard to calculation of methylation age where only smaller fractions seem to be apparent and to be unaffected by any potential proportional bias. One observation in our analyses relates to the difference in 7-CpG-derived DNAm age in men which on average was two years higher compared to women although the difference in chronological age between sexes in this dataset was only 0.2 years. This difference in DNAm age was reported for other versions of epigenetic clocks before [24, 25] and may reflect male-specific behavior (such as increased cigarette smoking or alcohol consumption) or may represent a bias due to other reasons and should be assessed further in independent datasets. Although cohorts that analyze the epigenetic clock [26-28] often contain older individuals whose mean chronological age is close to that of our participants, a more even age distribution would be desirable. Because the formula (“BII7”) that is used to calculate the 7-CpG clock was trained on the same participants but on samples taken on average 7.4 years earlier (at baseline examination), the individuals analyzed in this study are not independent from the 7-CpG training set. More data in independent datasets are needed to validate the conversion formula derived from our data.

In conclusion, we report a good degree of agreement between the individual methylation fractions of analyzed CpG sites measured with the SNuPE method and with the EPIC array by Illumina. A difference of 3.1 years between the DNAm age estimations based on the different measurement methods was found, which can be (partially) corrected by our newly developed conversion formula. With this study, we aim to increase the comparability between the 7-CpG clock determined with the SNuPE method and higher throughput methods, such as the EPIC array. Further studies are needed to clarify whether the high degree of agreement between methods can be replicated in different/younger age groups as well.

## Funding Statements

This work was supported by a grant of the Deutsche Forschungsgemeinschaft (grant number DE 842/7-1) to ID, the Cure Alzheimer’s Fund (as part of the “CIRCUITS-AD” consortium project) and the European Research Council’s “Horizon2020” funding scheme (as part of the “Lifebrain” consortium project; both to LB). This article uses data from the Berlin Aging Study II (BASE-II) and the GendAge study which were supported by the German Federal Ministry of Education and Research under grant numbers #01UW0808; #16SV5536K, #16SV5537, #16SV5538, #16SV5837, #01GL1716A and #01GL1716B.

## Acknowledgments

We thank Mrs. Sanaz Sedghpour Sabet and Tanja Wesse as well as Drs. Andre Franke and Michael Wittig for their help in generating the EPIC array data used in this study. We also acknowledge the high-performance compute environment (“OmicsCluster”) at University of Lübeck where the initial EPIC-data processing and analysis steps were run.

## Author contribution

Conceived and designed the study: ID and VMV. Contributed study specific data: all authors. Analyzed the data: VMV. Wrote the manuscript: VMV. All authors revised and approved the manuscript.

## Conflict of interest

None declared.

## Data availability statement

Data are available upon reasonable request. Interested investigators are invited to contact the study coordinating PI Ilja Demuth at to obtain additional information about the GendAge study and the data-sharing application form.

## Notes

### Competing Interest Statement

The authors have declared no competing interest.

