## Supplementary Material for "Seven-CpG DNA Methylation Age determined by Single Nucleotide Primer Extension and Illumina’s Infinium MethylationEPIC array provide highly comparable results"

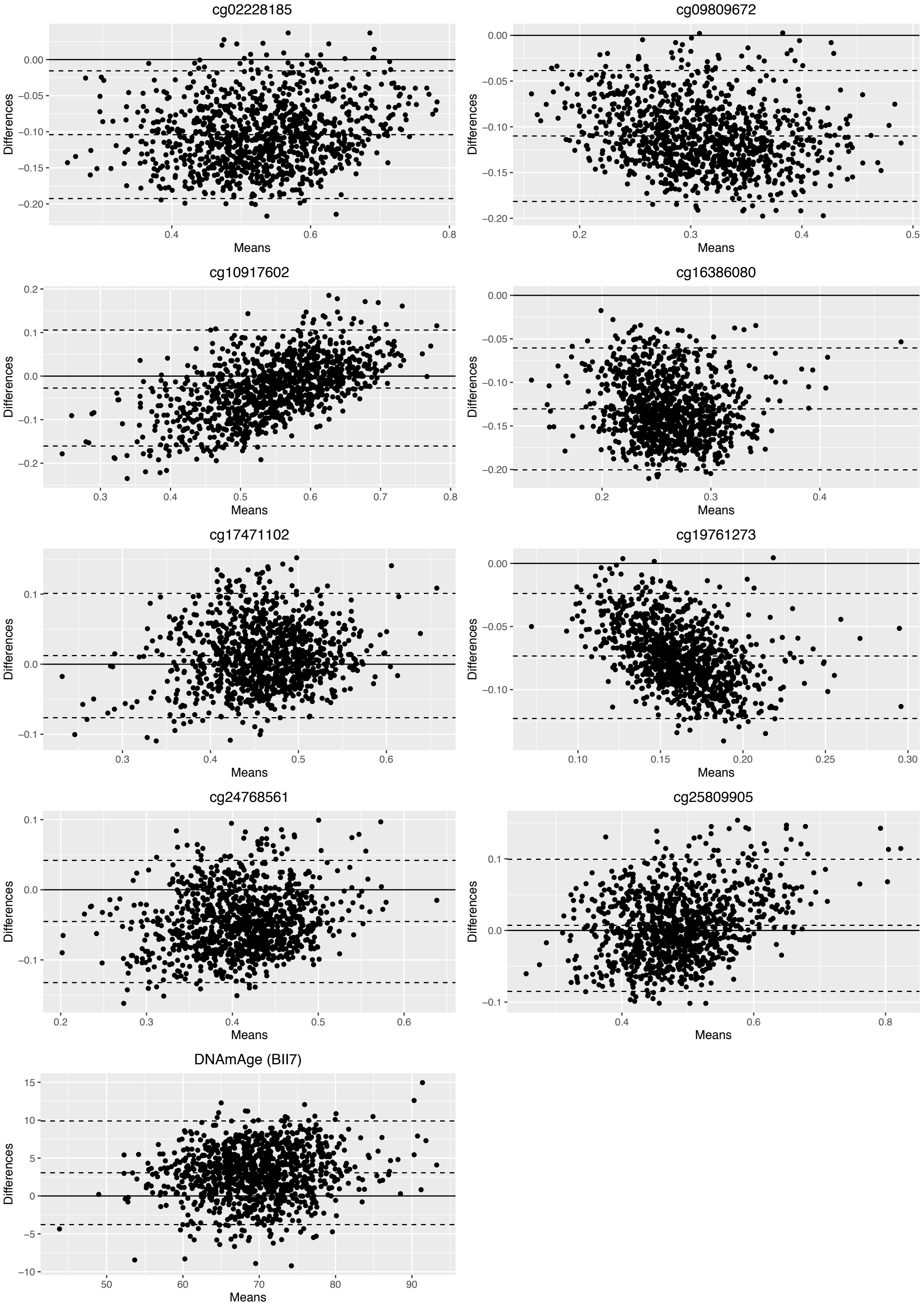


**Supplementary Figure 1:** Bland-Altman plots of the eight individual CpG sites and the resulting 7-CpG clock DNAm age.

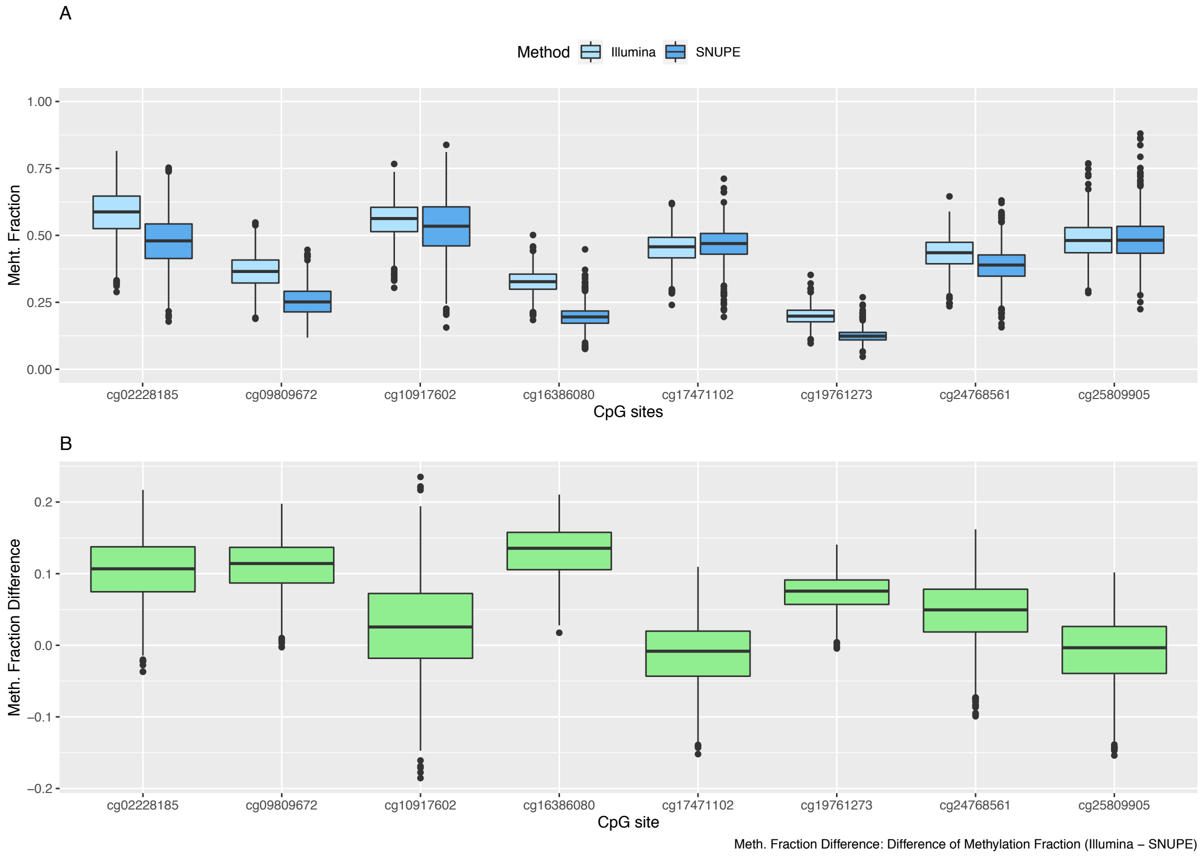


**Supplementary Figure 2:** Boxplots of methylation fraction measured by the SNuPE and EPIC array method (A) and difference between both methods (B). Cg10917602 was measured but is not used in the 7-CpG clock. In contrast to Figure 1 of the main manuscript it is included here as well.

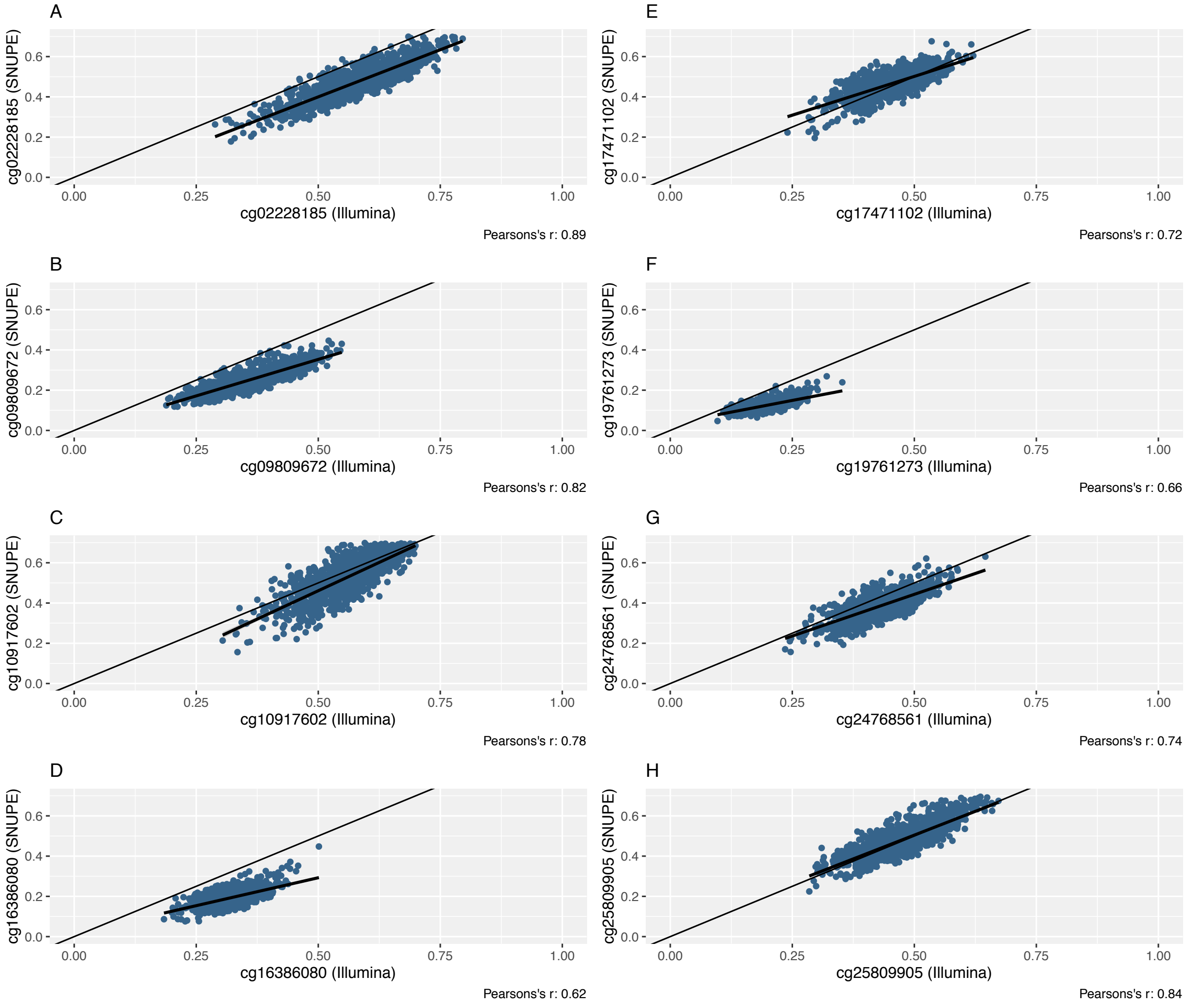


**Supplementary Figure 3:** Scatterplots of the methylation fraction of eight CpG sites that were measured with the SNuPE and EPIC array (Illumina) method (A-H). The line of equality (thin) and the regression line (bold) are displayed. In contrast to Figure 2 of the main manuscript, cg10917602 is included here as well.
